# Pro4S: prediction of protein solubility by fusing sequence, structure, and surface

**DOI:** 10.1101/2025.11.05.686869

**Authors:** Jie Qian, Lin Yang, Renxiao Wang, Yifei Qi

**Author notes:** Corresponding author Correspondence to: Renxiao Wang and Yifei Qi, Corresponding authors must provide their ORCID ID before resubmitting the final version of the manuscript. Non-corresponding authors are encouraged to link their ORCID. Contributions. J.Q., R.W. and Y.Q. conceived the research project. J.Q., L.Y. and Y.Q. designed and trained the model. J.Q. performed data analysis and wrote the paper. All authors read and commented on the paper.

## Abstract

Protein solubility is a critical physicochemical property influencing protein stability, therapeutic efficacy, and overall developability in drug discovery. However, traditional experimental methods for assessing solubility are often resource-intensive and time-consuming. To address these limitations, computational approaches leveraging artificial intelligence have emerged, yet current models generally treat qualitative classification and quantitative regression as separate tasks and rely predominantly on sequence-based information, neglecting crucial structural and surface characteristics. Here, we introduce Pro4S, a novel multimodal predictive model that integrates protein language models, structural data, and surface descriptors using advanced contrastive learning techniques. Our unified framework achieves significant improvements in prediction accuracy, robustness, and generalizability for both qualitative and quantitative solubility assessments. Benchmark comparisons demonstrate that Pro4S consistently outperforms existing state-of-the-art predictors across diverse datasets. Furthermore, by applying Pro4S to the emerging area of *de novo* protein design, we validated a strong correlation between predicted solubility and experimental expression levels, reducing the proportion of non-expressed proteins by 52.7% while retaining 96.7% of highly expressed proteins. This highlights Pro4S’s potential to serve as a reliable upfront screening tool for increasing expression success rates and accelerating rational protein engineering.

## Introduction

Proteins constitute the fundamental basis of life, participating extensively in physiological processes including cellular structural formation, biological catalysis, signal transduction, and immune responses.^1,2^ Their structural and functional diversity positions proteins as essential molecules for normal physiological activities, and also makes them critical targets in disease diagnosis, therapy, and drug discovery.^3–7^ In recent years, therapeutic proteins have become increasingly prevalent as macromolecular drugs, emphasizing the importance of deeply understanding and accurately predicting their physicochemical properties, such as solubility, aggregation propensity, expressibility, and immunogenicity.^8,9^

Protein solubility is crucial, directly influencing protein stability, interactions, and metabolic processes within cells.^10,11^ In drug discovery, particularly in evaluating therapeutic antibodies and enzyme-based pharmaceuticals, solubility serves as a key indicator of developability.^12^ Poor solubility often leads to instability, formulation challenges, and elevated immunogenicity risks. Moreover, for designed or engineered proteins, high solubility is critical to ensure successful expression and functionality. Accurate prediction of solubility is thus essential for improving the efficiency and success rate of biopharmaceutical development.^13,14^

However, current experimental methods for determining protein solubility generally require extensive time and human resources, involving complex processes such as protein expression, purification, and solubility assays.^15,16^ To address these challenges, computational approaches based on artificial intelligence have emerged in recent years, demonstrating significant application potential.^17–35^ Leveraging machine learning or deep learning algorithms, these methods utilize existing protein data to predict the solubility of unknown proteins automatically, significantly improving prediction efficiency and reducing experimental costs.^36^

In qualitative prediction, namely classification, most existing methods rely on sequence-based models such as DeepSoluE^37^, NetSolP^26^, Protein-Sol^31^, SKADE^38^, SoluProt^34^, and SWI^33^ (**Table 1**). These approaches typically employ protein language models, physicochemical amino acid features, or other descriptors to achieve accurate predictions. In the domain of quantitative prediction, namely regression, early sequence-based methods include DeepSol^19^, ProGAN^30^, SeqVec^39^, and TAPE^40^ (**Table 1**). With the advent of AlphaFold^41^, protein structure prediction has become more convenient and accurate, leading to the development of novel solubility prediction models that integrate structural information, such as GraphSol^35^, HybridGCN^23^, GATSol^22^, and ProtSolM^42^.

**Table 1.** Overview of protein solubility prediction models, their input modalities and training data types.

| Model | Modality |  |  | Training Data |  |
| --- | --- | --- | --- | --- | --- |
|  | Sequence | Structure | Surface | Qualitative | Quantitative |
| DeepSoluE<sup>37</sup> | √ |  |  | √ |  |
| NetSolP<sup>26</sup> | √ |  |  | √ |  |
| Protein-Sol<sup>31</sup> | √ |  |  | √ |  |
| SKADE<sup>38</sup> | √ |  |  | √ |  |
| SoluProt<sup>34</sup> | √ |  |  | √ |  |
| SWI<sup>33</sup> | √ |  |  | √ |  |
| ProtSolM<sup>42</sup> | √ | √ |  | √ |  |
| DeepSol<sup>19</sup> | √ |  |  |  | √ |
| ProGAN<sup>30</sup> | √ |  |  |  | √ |
| SeqVec<sup>39</sup> | √ |  |  |  | √ |
| TAPE<sup>40</sup> | √ |  |  |  | √ |
| GraphSol<sup>35</sup> | √ | √ |  |  | √ |
| HybridGCN<sup>23</sup> | √ | √ |  |  | √ |
| GATSol<sup>22</sup> | √ | √ |  |  | √ |
| Pro4S(Ours) | √ | √ | √ | √ | √ |

Despite considerable progress, existing prediction methods still exhibit certain limitations. Most studies currently separate qualitative prediction from quantitative prediction, lacking a unified prediction framework (**Table 1**).^22,26,27,34,43^ Furthermore, the majority of existing methods rely primarily on protein sequence information, inadequately accounting for crucial physicochemical features such as protein structure and surface properties, resulting in limited accuracy and generalizability.^19,26,30,33,38^ Previous research has indicated that protein solubility largely depends on spatial structure and surface characteristics.^36,44^ Structural properties such as surface hydrophobicity, polar residue distribution, and solvent-exposed surface area significantly influence protein solubility in aqueous environments.^36^ Consequently, relying solely on sequence information may be insufficient to fully capture these complex relationships.

To overcome these limitations, we propose Pro4S, a multimodal predictive model integrating protein sequence, structural, and surface property information (**Table 1**). By jointly modeling multiple protein features, Pro4S leverages advanced methods including ESM^45^, Masif^46^, AlphaFold^41^, and contrastive learning^47^. This approach aims to substantially enhance prediction accuracy and robustness, ultimately fulfilling practical demands in scientific research and pharmaceutical development.

## Results

### Overview of the Pro4S Model

Pro4S accepts Protein Data Bank (PDB) structures as input, extracts the amino-acid sequence, surface descriptors, and three-dimensional structural features, and integrates these representations to predict protein solubility (**Figure 1A**). Initially, the model extracts biochemical surface features and geometric structural features separately from the protein surface and its 3D backbone structure, respectively. These two sets of features are then projected linearly to the same dimension. The projected features are subsequently processed independently using Graph Neural Network (GNN) for iterative refinement.^48^ After each iteration, information is exchanged between the structural and surface graphs via a cross-graph interaction, enabling alternating fusion of the two graph representations (**Figure 1B**). A global node is explicitly maintained throughout this process to aggregate global information. This cycle is repeated multiple times to enhance the multimodal integration of features. To further improve consistency across modalities, a contrastive learning is introduced between the global node representation and the mean-pooled sequence embeddings, encouraging the encoder to capture shared information across modalities.

**Figure 1.**
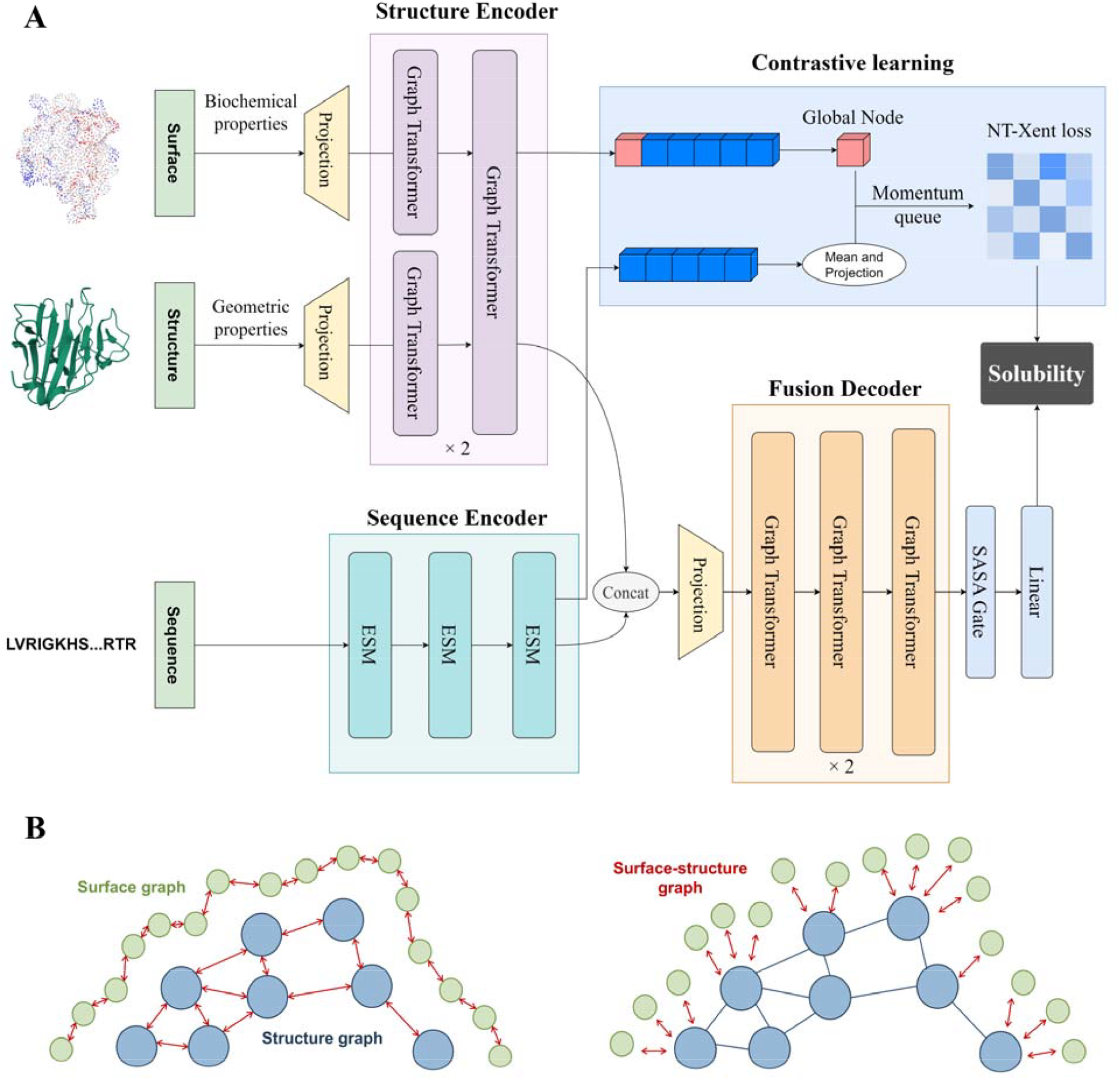
Overview of Pro4S. **A** The flowchart of the Pro4S framework. Surface biochemical features and backbone geometry are separately encoded and iteratively fused via cross-graph attention. A global node aggregates semantic information, guided by contrastive alignment with ESM-derived sequence embeddings. Fused features are decoded with SASA-gated GNN (Graph Neural Network) to predict solubility. **B** The structure encoder consists of three components: the surface graph, the structure graph, and the surface-structure interaction graph (detailed in Method). Information exchange occurs first within each component independently, followed by interaction between the surface and structural graphs.

**Figure 2.**
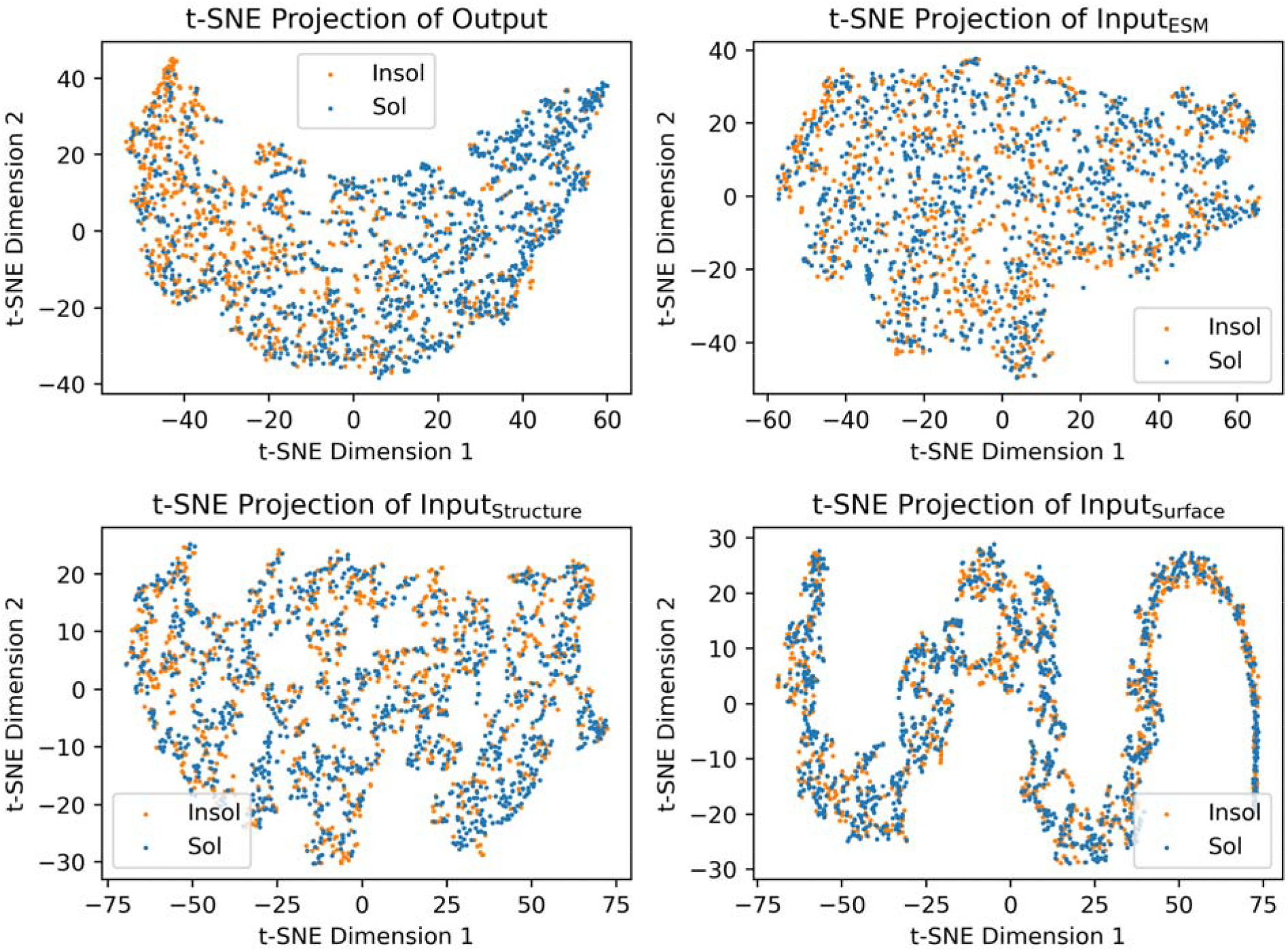
T-SNE visualization of the input and output features for Pro4S classification. The panels represent dimensionality reduction visualizations of the output layer (top left), ESM embeddings (top right), structural features (bottom left), and surface features (bottom right). Yellow points indicate insoluble proteins, while blue points indicate soluble proteins.

In parallel, sequence embeddings are generated from amino acid sequences using a protein language model ESM-3B.^45^ Structural representations and sequence embeddings are concatenated along the residue dimension and processed by a fusion decoder. This decoder, comprising multiple layers of GNNs, incorporates a linear projection at the input stage to ensure dimension alignment. To emphasize the relevance of solubility to surface residues, a solvent-accessible surface area (SASA)-based gating mechanism is incorporated at the decoder’s output, dynamically modulating channel features based on the residue-specific SASA values. Finally, the predictions are obtained by applying mean pooling across all residues, followed by a linear layer.

### Qualitative Classification of Protein Solubility

To evaluate the performance of our model in the qualitative task of solubility prediction, we employed the datasets from the previously published NetSolP study.^26^ In contrast to the original NetSolP study, which trained and tested on the PSI:Biology and Price datasets separately, we consolidated the these datasets for our training. PSI:Biology dataset contains 12,216 proteins expressed in *E. coli* using pET21 and pET15 vectors. The Price dataset contains 9,272 proteins that were expressed in *E. coli* using a unified production pipeline. The combined data, after preprocessing and removing redundant sequences with over 25% sequence identity, resulted in 10,671 proteins. The corresponding preprocessing steps are detailed in the Methods section. These were split into training and test sets in an 8:2 ratio, comprising 8585 and 2,086 sequences, respectively. We conducted a systematic performance comparison between our proposed model and several representative methods on the qualitative internal test set (**Table 2**). Our model consistently outperforms existing approaches across key metrics, achieving an AUC of 0.725, accuracy of 0.672, MCC of 0.334, and precision of 0.723. In particular, Pro4S achieved a 47.1% improvement in MCC over the best-performing baseline, SWI, highlighting its robustness and reliability. Moreover, the model maintains a high sensitivity while simultaneously improving specificity and F1-score, reflecting superior overall performance.

**Table 2.** Classification performance comparison of different models on the qualitative internal test set.

| Model<sup>a</sup> | AUC | Accuracy | F1-score | MCC | Precision | Recall | Specificity |
| --- | --- | --- | --- | --- | --- | --- | --- |
| ProtSolM | 0.511 | 0.577 | **0.722** | 0.050 | 0.580 | **0.957** | 0.065 |
| DeepSoluE | 0.654 | 0.613 | 0.657 | 0.213 | <u>0.668</u> | 0.646 | 0.568 |
| Protein-Sol | 0.641 | 0.598 | 0.643 | 0.183 | 0.655 | 0.632 | 0.552 |
| SKADE | 0.562 | 0.449 | 0.159 | 0.038 | 0.637 | 0.091 | **0.930** |
| SoluProt | 0.649 | 0.600 | 0.646 | 0.186 | 0.656 | 0.637 | 0.550 |
| SWI | <u>0.674</u> | <u>0.632</u> | <u>0.711</u> | <u>0.227</u> | 0.648 | <u>0.787</u> | 0.424 |
| Pro4S | **0.725** | **0.672** | 0.709 | **0.334** | **0.723** | 0.694 | <u>0.642</u> |

<sup>a</sup> Due to substantial overlap between this qualitative internal test set and the training data of the NetSolP model, NetSolP was not evaluated on this dataset.<sup>26</sup>

To intuitively illustrate the classification performance of Pro4S, we applied t-SNE dimensionality reduction to features extracted from the ESM embeddings, surface, structural characteristics, and the final classification layer outputs on the test dataset.^49^ Dimensionality reduction of raw ESM embeddings did not yield distinct clustering, suggesting that the model does not primarily rely on functional similarity or homologous information for solubility prediction. Conversely, features extracted from the classification layer effectively distinguished soluble from insoluble proteins, indicating robust classification performance.

To further validate the generalizability and robustness of our model in the binary classification task, we employed an external test set derived from a previous study.^27^ The qualitative external test set comprises 216 unpublished protein solubility data points clustered at 25% sequence identity, including tandem repeat proteins, DNA transposases, and deaminases. All test sequences share < 25% similarity with our training data, ensuring unbiased evaluation. As shown in **Table 3**, our model demonstrates a marked advantage in protein solubility prediction. Notably, our model achieves significant improvements of approximately 10 percentage points in both AUC and accuracy compared to state-of-the-art methods (**Figure 3A, Table 3**). Interestingly, the earlier machine learning model SWI maintained its superior performance across both test sets.^33^ This suggests that some deep learning models may overfit to their training data, limiting their effectiveness in real-world applications. These results indicate that our model has a stronger ability to differentiate between soluble and insoluble proteins and offers enhanced predictive robustness, underscoring its potential for practical use in solubility prediction.

**Table 3.** Classification performance comparison of different models on the qualitative external test set.

| Model | AUC | Accuracy |
| --- | --- | --- |
| ProtSolM | 0.50 | 0.38 |
| DeepSoluE | 0.62 | 0.54 |
| NetSolP | 0.68 | 0.49 |
| Protein-Sol | 0.67 | 0.57 |
| SKADE | 0.67 | <u>0.64</u> |
| SoluProt | 0.62 | 0.59 |
| SWI | <u>0.74</u> | 0.57 |
| Pro4S | **0.83** | **0.76** |

<sup>a</sup> For methods other than ProtSolM, performance values are adopted from the original PLM\_Sol paper.<sup>27</sup>

**Figure 3.**
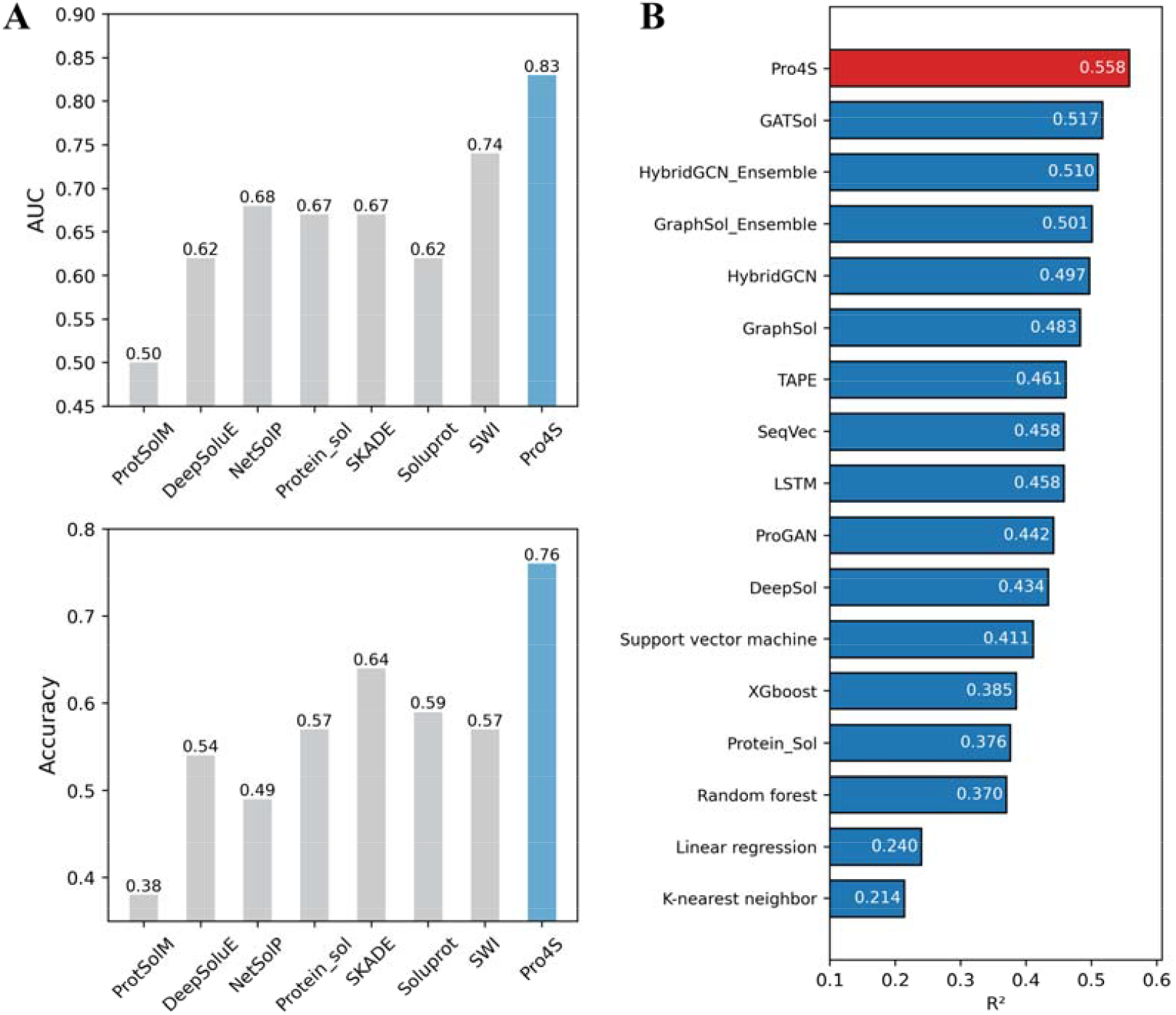
Performance of different models across test sets. **A** Classification performance of different models on the external test set (top: AUC, bottom: Accuracy). Data were sourced from a previous study.^27^ **B** Regression performance of different models on the internal test set. Data were sourced from a previous study.^22^

### Quantitative Regression of Protein Solubility

In the quantitative prediction of solubility, we employed the eSOL dataset^50^, following the same clustering procedures and dataset splitting strategy as established in previous work, ensuring comparability across experiments.^22,35^ The eSOL database provides comprehensive solubility data for the *E. coli* proteins, obtained using the PURE system without chaperones.^50^ By benchmarking the proposed model against several mainstream methods for protein solubility prediction, our model showed significant improvements across key performance metrics. As shown in **Tables 4** and **Figure 3B**, conventional machine learning methods such as Random Forest (R^2^ = 0.370) and XGBoost (R^2^ = 0.385) exhibited limited predictive capabilities. Deep learning approaches, including DeepSol (R^2^ = 0.434) and TAPE (R^2^ = 0.461), demonstrated moderate improvement yet still left room for further enhancement. Our model achieved an R^2^ of 0.558, representing a significant improvement over the current best-performing model, GATSol (R^2^ = 0.517). This breakthrough demonstrates the enhanced capability of our new model in accurately predicting protein solubility. Furthermore, our approach outperformed existing methods across multiple metrics, including accuracy (0.808), F1-score (0.783), and AUC (0.896), underscoring the effectiveness of our innovative model architecture. The substantial improvement in R^2^ directly reflects the superior predictive power of the proposed multimodal method, highlighting its potential applicability in protein engineering tasks.

**Table 4.**
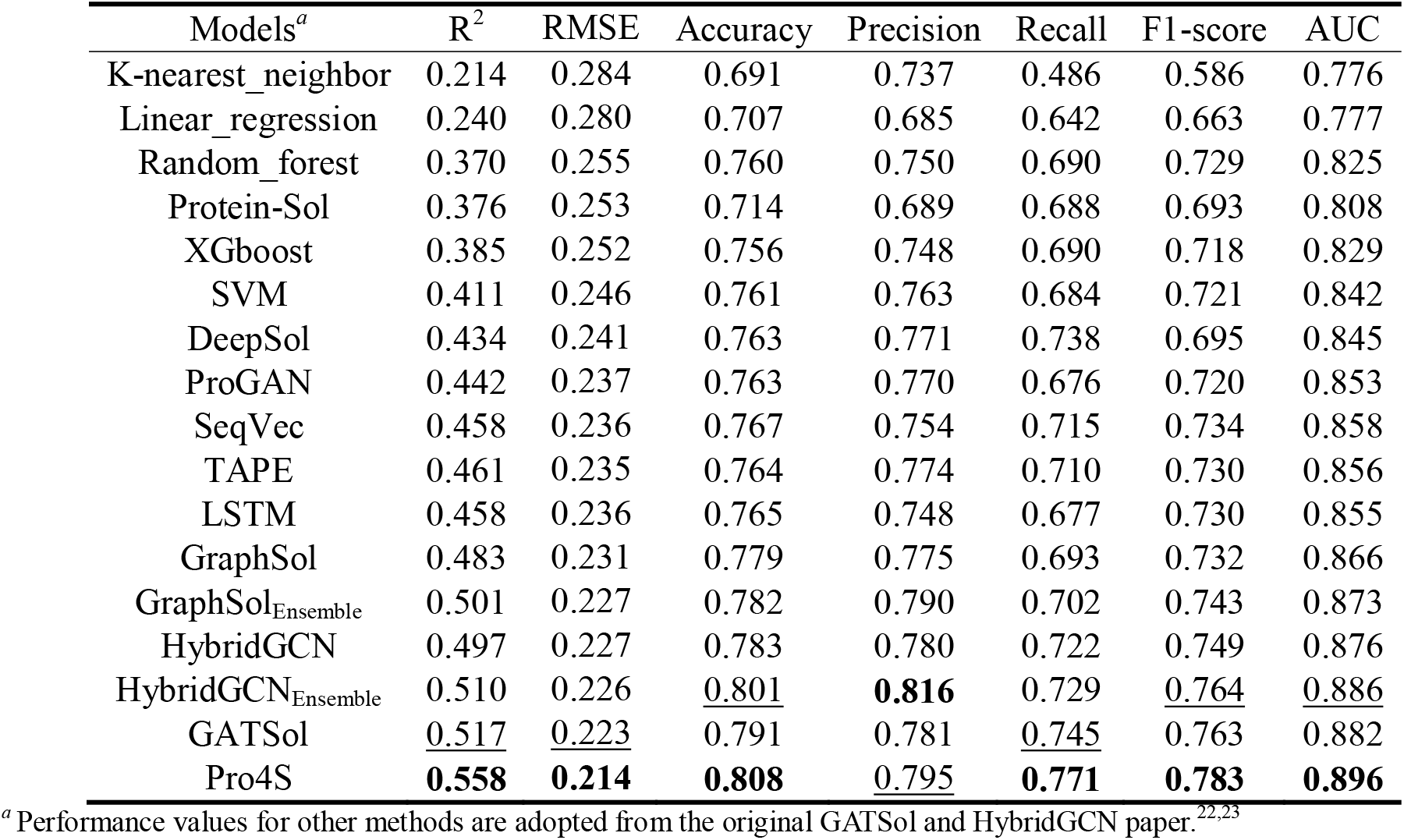
Regression performance comparison of different models on the quantitative internal test set (N = 660).

To further validate the generalizability and robustness of our model in the quantitative prediction task, we employed an external test set derived from a previous study.^35^ The quantitative external test set comprised 108 *Saccharomyces cerevisiae* proteins with corresponding experimental structures. Solubility values for this set were experimentally determined using a cell-free expression system (PURE). As summarized in **Table 5**, methods such as ProGAN (R^2^ = 0.08) and DeepSol (R^2^ = 0.09) exhibited limited predictive accuracy. By contrast, graph neural network-based models demonstrated clear advantages, with GraphSOL_Ensemble_ and HybridGCN both achieving an R^2^ of 0.37, and HybridGCN_Ensemble_ further improving to 0.39. Particularly noteworthy is the GATSol model, which significantly improved prediction performance (R^2^ = 0.42) by incorporating graph attention mechanisms. Our proposed model delivered the highest performance (R^2^ = 0.43), surpassing the current best methods. These findings validate the effectiveness of our model’s architectural enhancements and suggest that integrating surface features alongside structural information substantially improves prediction accuracy.

**Table 5.** Regression performance comparison of different models on the quantitative external test set (N = 108)

| Model<sup>a</sup> | $R^2$ |
| --- | --- |
| ProGAN | 0.08 |
| DeepSol | 0.09 |
| Protein-Sol | 0.28 |
| ccSol | 0.30 |
| GraphSOL | 0.36 |
| GraphSOL<sub>Ensemble</sub> | 0.37 |
| HybridGCN | 0.37 |
| HybridGCN<sub>Ensemble</sub> | 0.39 |
| GATSol | <u>0.42</u> |
| Pro4S | **0.43** |

<sup>a</sup> Performance values for other methods are adopted from the original GATSol and HybridGCN paper.<sup>22,23</sup>

### *De novo* protein design test set

Protein solubility prediction can reflect protein expression levels to a certain extent. To evaluate the generalization ability of solubility prediction models in *de novo* designed protein expression tasks, we extracted protein expression data from the recent Adaptyv EGFR Protein Design Competition as an external test set.^51^ Following rigorous data cleaning procedures (details provided in the Methods), we obtained 147 valid data points, comprising 55 proteins with very low expression and 99 proteins with high expression. This competition have indicated that solubility prediction models such as SoluProt and NetSolP can enhance protein expression rates.^51^ Therefore, we included these two models along with other leading solubility prediction models as baselines. Furthermore, this work has identified the proportions of GLU and LYS residues in designed proteins as robust predictors of solubility.^51^ Consequently, we incorporated these residue proportions as baselines of protein expression in our analysis.

We systematically compared three Pro4S variants (Pro4S_classification_, Pro4S_regression_, and Pro4S_finetune_) against mainstream existing models. The Pro4S_classification_ and Pro4S_regression_ models, as previously mentioned, were trained on classification and regression data, respectively. Building upon the Pro4S_classification_ model, we developed the Pro4S_finetune_ model through further training on the quantitative eSOL dataset. As shown in **Tables 6**, all three variants demonstrate notable advantages across multiple core metrics: Pro4S_finetune_ achieved the best overall performance with an accuracy of 0.803, an F1-score of 0.860, and an MCC of 0.580, along with approximately a 42% improvement in specificity (0.527) compared to baseline averages. Pro4S_regression_ delivered the highest performance in terms of AUC (0.784), outperforming the second-best baseline model by 8.4%. Pro4S_classification_ displayed remarkable balance in precision (0.769) and specificity (0.545). Of particular note, all three Pro4S variants showed significant improvements in MCC values (0.491-0.580), demonstrating superior predictive performance compared to existing methods, regardless of training on classification or regression data. These results underscore the breakthrough capability of the Pro4S framework in predicting the expressibility of *de novo* designed proteins.

**Table 6.** Performance comparison of different models on *de novo* protein design test set (N = 147).

| Model | AUC | Accuracy | F1-score | MCC | Precision | Recall | Specificity |
| --- | --- | --- | --- | --- | --- | --- | --- |
| ProtSolM | 0.500 | 0.626 | 0.770 | 0.000 | 0.626 | **1.000** | 0.000 |
| SKADE | 0.543 | 0.580 | 0.677 | 0.083 | 0.643 | 0.716 | 0.364 |
| DeepSoluE | 0.582 | 0.604 | 0.721 | 0.102 | 0.628 | 0.845 | 0.236 |
| LYS_ratio | 0.712 | 0.633 | 0.722 | 0.188 | 0.686 | 0.761 | 0.418 |
| NetSolP | 0.689 | 0.639 | 0.773 | 0.125 | 0.638 | 0.978 | 0.073 |
| SoluProt | 0.637 | 0.664 | 0.776 | 0.243 | 0.659 | 0.943 | 0.218 |
| Protein-Sol | 0.730 | 0.686 | 0.790 | 0.336 | 0.664 | 0.976 | 0.236 |
| SWI | 0.703 | 0.693 | 0.749 | 0.354 | 0.744 | 0.753 | **0.600** |
| GLU_ratio | 0.720 | 0.694 | 0.759 | 0.339 | 0.747 | 0.772 | 0.564 |
| GATSol | 0.723 | 0.755 | 0.830 | 0.468 | 0.733 | 0.957 | 0.418 |
| Pro4S<sub>classification</sub> | 0.762 | 0.769 | 0.830 | 0.491 | <u>0.769</u> | 0.902 | <u>0.545</u> |
| Pro4S<sub>regression</sub> | **0.784** | <u>0.776</u> | <u>0.847</u> | <u>0.533</u> | 0.740 | <u>0.989</u> | 0.418 |
| Pro4S<sub>finetune</sub> | <u>0.762</u> | **0.803** | **0.860** | **0.580** | **0.774** | 0.967 | 0.527 |

Analysis of the confusion matrix for the fine-tuned Pro4S model revealed that protein screening with this model significantly reduces the proportion of inexpressible proteins by approximately 50% while maintaining nearly constant recovery of expressible proteins (**Figure 4A**). Consequently, the overall protein expression rate improved from 67.3% to 77.4%.

**Figure 4.**
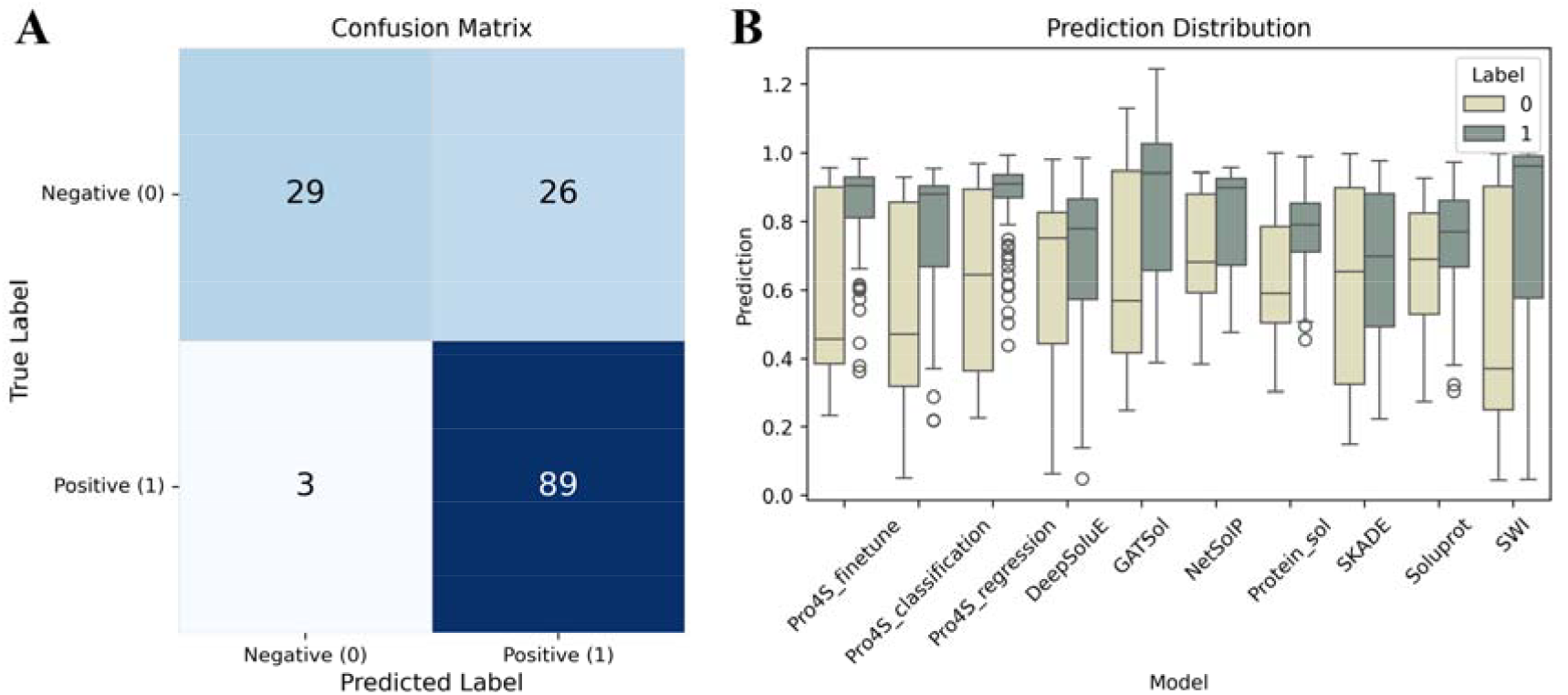
Evaluation on *de novo* protein design test set. **A** Confusion matrix illustrating the prediction performance of the Pro4S_finetuned_ model. **B** Comparative distributions of predicted scores for expressible and inexpressible proteins by various models evaluated on the *de novo* protein design test set. ProtSolM model is not shown because its output labels were all 1.^42^

To visualize the distribution patterns of prediction labels across different models, we compared the output scores of Pro4S variants and baseline methods as illustrated in **Figure 4B**. The Pro4S variants (Pro4S_finetune_, Pro4S_classification_, and Pro4S_regression_) consistently demonstrated superior discriminatory power compared to existing methods. Moreover, predictions for positive samples (expressible proteins; Label=1) were highly concentrated within a narrow interval, indicating high and stable confidence in the model’s predictions of expressible sequences. For negative samples (inexpressible proteins; Label=0), the median prediction scores were effectively suppressed to between 0.30 and 0.45, and the distribution tails substantially reduced, highlighting the model’s robust ability to identify inexpressible sequences. In contrast, representative baseline models such as DeepSolE, GATSol, and NetSolP exhibited significant overlap in their positive and negative sample distributions, with median prediction scores for positives frequently below 0.85 and for negatives typically above 0.55. This overlap implies greater predictive bias and broader error margins near decision boundaries. Additionally, the machine learning-based method SWI acts as “a stringent filter”, excluding a higher proportion of non-expressed proteins (SWI 60.0% vs. Pro4S_finetune_ 52.7%) but at the cost of discarding more expressed proteins (SWI 24.7% vs. Pro4S_finetune_ 3.3%). Notably, Pro4S_finetune_ combines the strengths of the regression model’s ability to precisely identify positive samples and the classification model’s efficacy in suppressing negative samples, further demonstrating that fine-tuned Pro4S can effectively minimize misclassification while maintaining high confidence. Overall, the Pro4S series achieve a median prediction gap exceeding 0.40 between positive and negative samples, considerably broadening the decision boundary and providing significantly improved reliability in probabilistic outputs compared to the best available baseline models. This substantial enhancement facilitates more accurate and efficient high-throughput protein expressibility screening.

### Ablation experiment

In the ablation study, we systematically evaluated the contribution of each component in the Pro4S model. The full model achieved optimal performance in both qualitative (AUC = 0.725) and quantitative (R^2^ = 0.558) tasks (**Figure 5**). Upon sequential removal of key modules, we observed a stepwise decrease in model performance. Removing the contrastive loss led to a reduction in AUC and R^2^ to 0.720 and 0.550, respectively, indicating that contrastive constraints effectively enhance the discriminative boundaries and prediction accuracy. Further exclusion of surface property features caused the performance to decrease to AUC = 0.714 and R^2^ = 0.535, demonstrating the critical role of surface physicochemical information in capturing functional signals. Concurrent removal of structural and surface features resulted in an even more pronounced performance drop (AUC = 0.708, R^2^ = 0.528), highlighting the synergistic effects of structural and surface information. Finally, when the ESM language model embedding was entirely omitted, the model exhibited the lowest performance (AUC = 0.685, R^2^ = 0.420), underscoring the fundamental impact of deep sequence representations on overall predictive capability. Taken together, each module contributes distinctly to the predictive power of the model, with ESM embeddings providing the core informational foundation, while structural-surface features and contrastive loss further enhance performance through complementary spatial and physicochemical cues.

**Figure 5.**
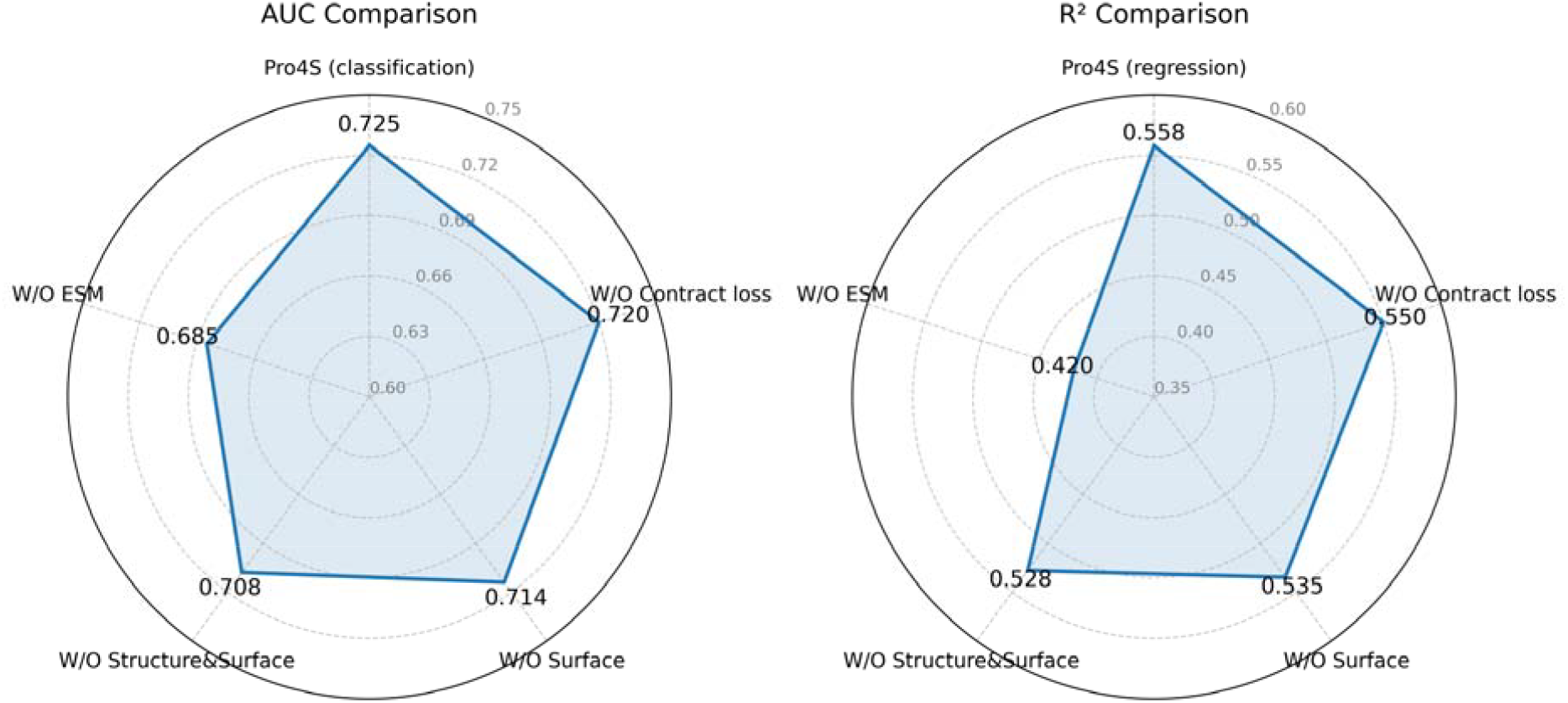
Ablation analysis of the Pro4S framework. Left, radar plot of area under the ROC curve (AUC) for the classification task; right, radar plot of coefficient of determination (R^2^) for the regression task. Each vertex of the radar chart represents the result after ablating a certain component in the complete Pro4S model.

## Conclusion

We present a novel and efficient protein solubility prediction model that integrates protein language models, structural features, and surface descriptors into a unified multimodal framework, substantially improving the accuracy and reliability of solubility estimation. Across both qualitative classification and quantitative regression tasks, our approach consistently outperforms conventional solubility predictors, regardless of the training data type, demonstrating outstanding generalization and robustness.

Beyond benchmarking, we successfully applied the model to the burgeoning field of *de novo* protein design. Using proteins from the EGFR design challenge as a case study, we uncovered a strong correlation between predicted solubility and experimental expression levels, indicating that solubility prediction can be effectively extended to forecasting protein expression yields. Our model achieves state-of-the-art performance in this regard. Consequently, incorporating our predictor as an upfront screening tool is expected to markedly increase the expression success rate during protein design cycles, thereby accelerating *de novo* protein engineering. This work not only provides a powerful computational aid for protein engineering, but also opens new avenues for rational protein design.

## Methods

### Data processing

To ensure consistency and comparability with established methods, the data processing workflows for the two tasks (classification and regression) were conducted in accordance with the procedures outlined in prior studies.^22,27^ For the qualitative classification model, we adopted the data splitting procedure from PLM_Sol^27^, while for the quantitative regression model, we employed the protocol established in the GATSol^22^ work.

#### Classification task

We combined two high-quality solubility datasets from the NetSolP study—the PSI:Biology dataset and the Price dataset—for model training.^26^ To maximize representativeness and diversity, we adopted the data partitioning protocol established in prior work.^27^ Specifically, we first merge the three subsets and then use MMseqs2 clustering with a 75% sequence-identity cutoff to remove redundancy, selecting one sequence per cluster.^52^ The resulting sequences are split into training, validation, and test sets at a 8:2 ratio, with inter-set identity capped at 25%. This yields 10,671 sequences in total with 8585 for training and 2,086 for testing.

#### Regression task

We adopt the standard eSOL dataset. Sequences are deduplicated at a 30% identity threshold, following established practice^22^:

1. Prepare a dataset in FASTA format for homologation.
2. Extract the first amino acid sequence from the dataset to initialize a library file.
3. Retrieve the second protein sequence from the dataset and compare homology with the library file using ggsearch36.^53^ The sequence is added to the library file if the global identity between the two sequences is < 30% and the E-value is ≤ 1×10^−6^.
4. Repeat the above step for the third protein sequence and continue until all data in the dataset have been homologously compared once.

The final homology-separated dataset comprises 2679 protein sequences, with 2019 sequences (75%) allocated to the training set and 660 (25%) to the test set.

#### *De novo* protein design test set

To further evaluate different models in a protein-design scenario, we used protein-expression data from the first-round of Adaptyv EGFR *de novo* protein design competition. The competition categorizes expression levels into None, Low, Medium, and High. We label “None” as 0 and “High” as 1, retaining sequences whose triplicate measurements are consistently “None” or “High” to minimize stochastic noise. The resulting dataset contains 147 sequences—55 with negligible expression (label 0) and 92 with high expression (label 1).

### Graph representation

#### Structure graph

Since the training data consist of sequences paired with corresponding solubility labels, all protein structures were predicted using AlphaFold2.^41^

##### Node features

Each amino-acid node is characterized by a feature vector ***n***_*i*_ ∈ ℝ^36^, that integrates sequence, physicochemical and structural descriptors. The feature comprises: (i) a 20-dimensional one-hot representation of the residue type; (ii) physicochemical attributes: including hydrophobicity, charge, polarity and molecular mass; (iii) secondary-structure class (helix, strand or coil); (iv) solvent-accessible surface area (SASA); (v) per-residue confidence from structure prediction (pLDDT); and (vi) a binary flag indicating whether the residue is surface-exposed. We performed a nearest-neighbor search for each surface point, labeling the closest structural point to each surface point as 1, while all other points were labeled as 0.

##### Edge features

Prior studies have indicated that varying cutoff values have minimal impact on the results.^22,54^ We calculated the *C*_*β*_ distances between different residues to establish edges, with residue pairs set at 10 Å cutoff. For glycine, which lacks *C*_*β*_ atoms, we used the MP-NeRF method to compute the *C*_*β*_ atom coordinates.

For the interaction edges between amino-acid nodes, we constructed an equivariant edge feature vector comprising the following elements.

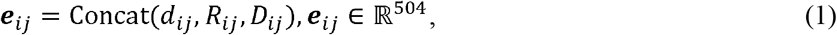

where *d*_*ij*_ is the distance feature, *R*_*ij*_ is the angle feature, and *D*_*ij*_ is the direction feature (described in detail below).

Distance features were computed between the two sets of atom coordinates, A and B:

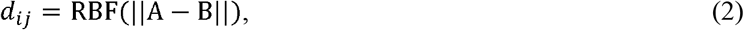

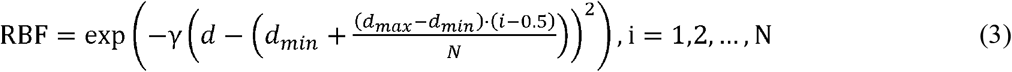

where A ∈ {*N*_*i*_, *C*_*ai*_, *C*_*i*_, *O*_*i*_, C _*βi*_ …},B ∈ {*N*_*j*_,*C*_*aj*_, *C*_*j*_, *O*_*j*_, *C* _*βi*_ …}. RBF represents the radial basis function. *d*_*min*_ and *d*_*max*_ define the range of distances over which the RBF function operates. Here, the default range is set from 0 to 20. The parameter governing the number of Gaussian basis functions (N) employed in the radial basis function expansion determined their spatial distribution within this interval. In the current implementation, a set of 16 uniformly distributed kernels is automatically generated across the predefined interval. We computed distances between all pairs of atoms in the two residues. In order to ensure that different residue pairs form edges of the same dimension, we selected distances between the five backbone atoms in each residue, resulting in a total of 25 distances. However, since backbone atoms may not interact, we included the five shortest distances between all atoms in residues *i* and *j* to capture potential interactions, resulting in a total of 30 distances. Eventually, we obtained the distance characterization *d*_*ij*_ ∈ ℝ^(3 ×16)^.

The angle features were characterized by a rotation matrix *R*_*ij*_ was obtained by multiplying the rotation matrices of the two residues corresponding to the edge.

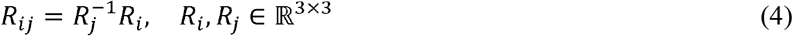

Here, we obtained rotation matrix *R*_*i*_,*R*_*j*_ based on the Gram-Schmidt orthogonalization process.^56^

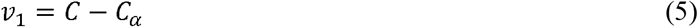

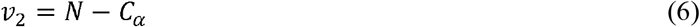

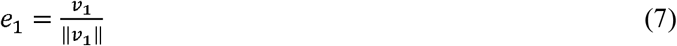

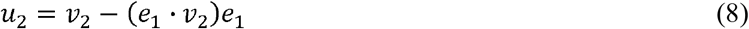

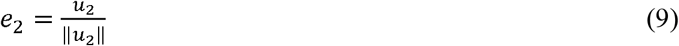

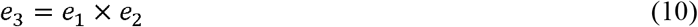

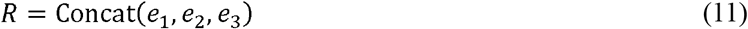

Direction features *D*_*ij*_ were defined using unit vectors between the *C* _*α*_ of the starting residue and the five backbone atoms of the target residue. We multiply this vector by 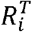 (obtained from angle features) to ensure its rotational invariance.

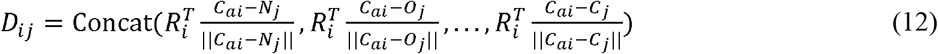

#### Surface graph

##### Node features

For each protein, solvent-exposed surface features were computed with the MaSIF, which renders the surface as a point cloud. Each surface point was represented as a graph node, characterized by a feature vector ***n***_*i*_ ∈ ℝ^7^, comprising hydrophobicity, hydrogen-bond donor/acceptor propensity, atomic charge, the local electrostatic field vector, and the Euclidean distance to the protein centroid. To preserve SE(3)-equivariance, a local reference frame was derived from principal-component analysis of the point cloud: (i) the geometric centre of the points served as the origin; (ii) the covariance matrix of point coordinates was diagonalised; (iii) the eigenvectors were ordered by decreasing eigenvalue and assigned as the X, Y and Z axes of a right-handed frame. Each surface normal was then expressed in this basis, enriching the geometric description. Because raw point clouds are dense, an octree-based down-sampling scheme was applied to eliminate redundant points and accelerate subsequent calculations.^57^ The octree root encompassed the point cloud’s bounding box, with center as the midpoint of min/max coordinates and size as the maximum axis extent. Construction recursed to a fixed max depth of 6, subdividing nodes with over an initial threshold of 20 points per node. To reach a target reduction to ∼1/20th of original points, we iteratively tuned this threshold (up to 5 iterations, or 1 for clouds <30,000 points). A comparison of point distributions before and after compression is provided in **Figure 6**.

**Figure 6.**
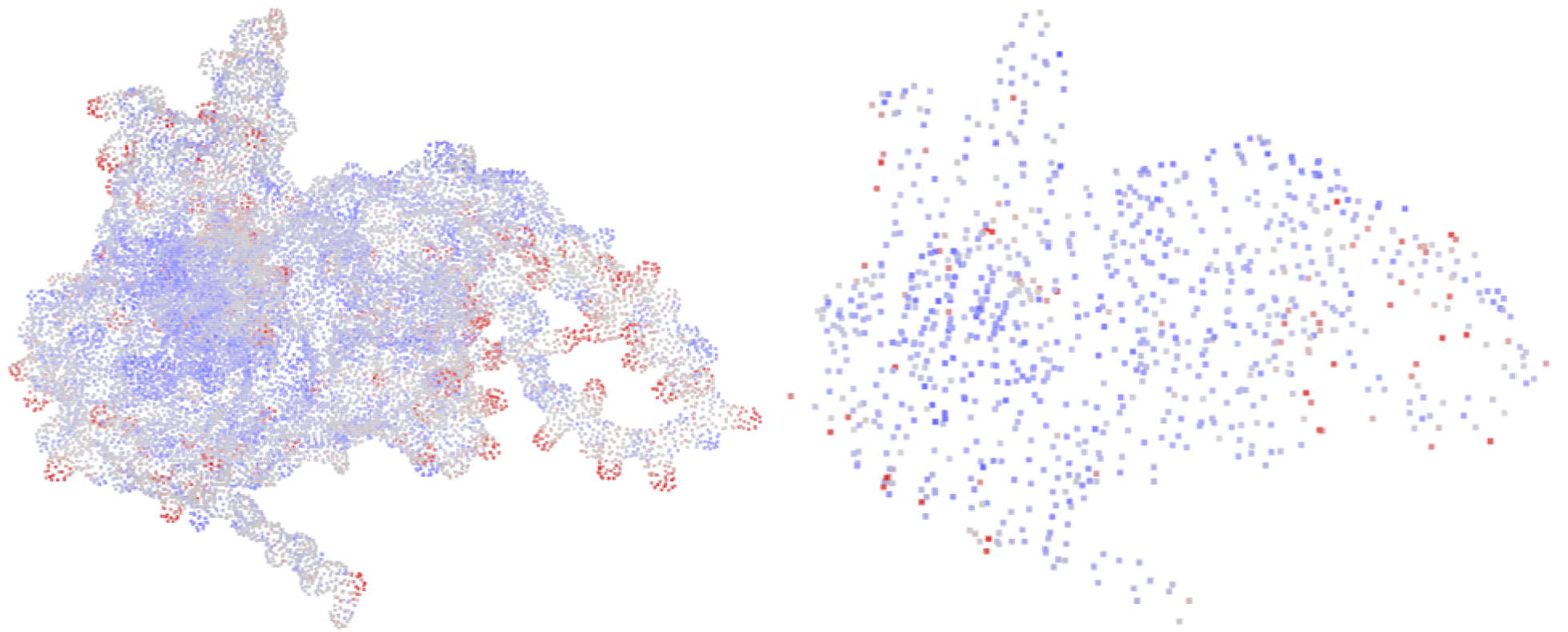
Protein surface generation. Left, raw surface. Right, compressed surface. Positively and negatively charged points are colored red and blue, respectively.

##### Edge features

For each pair of surface nodes A and B, the edge features were defined by applying a radial-basis function to the Euclidean separation of each pair of, yielding *d*_*ij*_ RBF (||A − B||).

#### Structure-surface graph

##### Node features

We integrated structural and surface graphs, with node features derived from surface and structural graphs updated via graph neural network.

##### Edge features

For each surface node, edge was established by connecting it to the nearest amino acid (**Figure 1B**). For each pair of surface node A and structure node B, the edge features were defined by applying a radial-basis function to the Euclidean distance of each pair, yielding *d*_*ij*_ RBF (||A − B||).

### Model training

#### Contrastive learning

For each protein structure, positive pairs are formed by associating its global structural aggregator node with the averaged sequence features. Negative pairs are constructed using two dynamically updated queues, Q_T_ and Q_S_, which store global structural embeddings and sequence embeddings, respectively, from the most recent 64 proteins encountered during training. The contrastive loss is defined as:

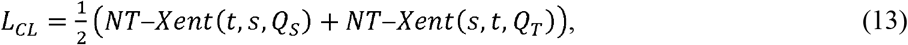

where NT-Xent is the normalized temperature-scaled cross entropy loss function, a contrastive loss used to maximize agreement between similar pairs of embeddings and push dissimilar pairs apart.^47^ s and t denote the global structure embeddings and sequence embeddings, respectively. The model is trained using a combined loss function: L = L_CE_ + L_CL_, where L_CE_ is the cross-entropy loss for solubility prediction.

#### Training objectives

In this study, we conducted five-fold cross-validation on both the qualitative and quantitative training datasets.^58^ Each dataset was randomly partitioned into five subsets. During each iteration, four subsets were used for model training, and the remaining subset was reserved for evaluation. This procedure was repeated five times, with hyperparameter optimization guided by the average AUROC (for qualitative tasks) and R^2^ (for quantitative tasks) across all rounds. After identifying the optimal hyperparameters, the model was retrained on the full training set and independently evaluated on separate test datasets.

We systematically evaluated both classification and regression tasks, ultimately determining unified model parameters. The hidden dimension of all GNN was set to 256, with both the structure encoder and fusion decoder configured to two layers. Training efficiency was optimized through an early-stopping strategy based on validation set performance, terminating training if no improvement was observed after two epochs. A dropout layer (with a rate of 0.5) is applied subsequent to the attention weight calculation to mitigate model overfitting. Concurrently, a Batch Normalization layer has been incorporated into the edge feature projection module. For both classification and regression tasks, all trainable parameters within our custom graph neural network are initialized via the Xavier uniform distribution, culminating in a total model size of approximately 15 M parameters, excluding the ESM parameters. We employed the Adam optimizer with a learning rate of 3×10^−5^.^59^ Models were trained to minimize prediction error using a batch size of 1. The loss function combined binary cross-entropy (BCE) and the NT-Xent contrastive loss. Training was conducted on an NVIDIA GeForce RTX 3090 GPU.

## Data Availability

The sequences of the training and test sets, along with their corresponding AlphaFold2-predicted structures, are available at our GitHub repository: https://github.com/TEKHOO/Pro4S.

## Code Availability

Installable source code, associated guidelines, various custom scripts, and interactive data analysis notebooks are available at GitHub https://github.com/TEKHOO/Pro4S.

## Acknowledgements

This study was financially supported by the National Natural Science Foundation of China (Grants No. 22373020 and 22033001 to Y. Qi, and No. 22537001 and 82173739 to R. Wang), the Shanghai Municipal Science and Technology Commission (Grant No. 25JS2830200 to R. Wang), and AI for Science Foundation of Fudan University (FudanX24AI050). The computations in this research were performed using the CFFF platform of Fudan University.

